# A faster implementation of association mapping from k-mers

**DOI:** 10.1101/2020.04.14.040675

**Authors:** Zakaria Mehrab, Jaiaid Mobin, Ibrahim Asadullah Tahmid, Atif Rahman

## Abstract

Genome wide association studies (GWAS) attempt to map genotypes to phenotypes in organisms. This is typically performed by genotyping individuals using microarray or by aligning whole genome sequencing reads to a reference genome. Both approaches require knowledge of a reference genome which limits their application to organisms with no or incomplete reference genomes. This caveat can be removed using alignment-free association mapping methods based on k-mers from sequencing reads. Here we present an implementation of an alignment free association mapping method [1] to improve its execution time and flexibility. We have tested our implementation on an *E. Coli* ampicillin resistance dataset and observe improvement in performance over the original implementation while maintaining accuracy in results. Finally, we demonstrate that the method can be applied to find sex specific sequences.

## Introduction

Association mapping is the process of associating phenotypes with genotypes. In genome wide association studies (GWAS), individuals are typically genotyped using mircoarrays or by aligning sequencing reads from individuals to a reference genome. However, both these approaches require a reference genome of the organism which makes them inappropriate for association mapping in non-model organisms with incomplete reference genome or none at all.

To address this issue, alignment free approaches for association mapping have been explored. A number of methods have been developed for association mapping in bacterial genomes since the high plasticity in those genomes makes application of reference based methods difficult [2, 3, 4, 5]. Rahman et al. [1] and Voichek et al. [6] presented methods for mapping associations in large genomes, to categorical phenotypes and to both categorical and quantitative phenotypes, respectively. The methods are primarily based on finding k-mers i.e. contiguous sequences of length k in sequenced reads and identifying k-mers associated with the phenotype.

In the association mapping tool named Hawk developed by Rahman et al. [1], frequencies of k-mers are analyzed to find k-mers associated with a phenotype and then they are assembled to form the associated sequences. First, they count k-mers in reads from each individual using Jellyfish [7]. Second, using likelihood ratio test, they find k-mers with significantly different counts in case and control samples. Next, population structure is determined from k-mer counts using Eigenstrat [8, 9]. After that, association to k-mers after correcting for population structure is determined. Finally, the k-mers found associated may be assembled to get a sequence for each associated loci.

Here we re-implement Hawk with the goal to reduce its execution time and make it more convenient for users. We have re-implemented the step for finding associated k-mers after population structure correction using C++, which was previously implemented in R. We have also extended support for Jellyfish 2 and implemented Benjamini–Hochberg procedure [10], which can be used to correct for multiple tests when the study is underpowered for Bonferroni correction. We have tested our implementation with a dataset on *E*.*coli* ampicillin resistance and have compared its output with the output of the original implementation. We have also analyzed the execution times of the two implementations. Our implementation is faster and more flexible to run compared to the original implementation while maintaining accuracy.

Finally, we show that our method can be used to find sequences in the sex chromosomes. We apply our method to sequencing data from two populations in the 1000 genomes dataset, labeling males and females as cases and controls. We find the k-mers determined by Hawk cover the entire sequenced regions in X and Y chromosomes. It is worth noting that other reference free methods for association mapping mentioned above are based on presence and absence of k-mers and hence are not suitable for finding sequences in sex chromosomes present in both sexes e.g. the human chromosome X.

## Improvement and New Features

Here we summarize the improvements and new features we have added. Results supporting the improvements is presented in the following section.

### Re-implementation of correction for population structure

Population stratification is a known confounder in association studies. Without correcting for this confounding factor, one may falsely associate non-significant genotypes with phenotypes. In the Hawk pipeline, population structure was estimated using Eigenstrat [8, 9] and subsequently p-values were adjusted for population structure using the glm function (for fitting logistic regression models) and the ANOVA function (for testing the goodness of fit) in R. We re-implement this process in C++ and thereby improving the performance.

R uses the IRLS (Iteratively Re-weighted Least Square) method to fit the model [11]. Therefore in our implementation of the glm function in C++, we also used IRLS for fitting the model. Iteratively re-weighted least squares for finding the MLE (Maximum Likelihood Estimate) for logistic regression is a special case of Newton’s algorithm. If the problem is written in vector matrix form, with parameters *w*^*T*^ = [*β*_0_, *β*_1_, *β*_2_,…], explanatory variables *x*(*i*) = [1, *x*_1_ (*i*), *x*_2_ (*i*),…]^*T*^ and expected value of Bernoulli distribution 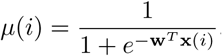, the parameters **w** can be found using the following iterative algorithm:

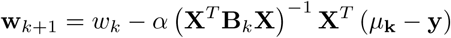

where *α* is the learning rate, **B** = diag(*µ*(*i*)(1 − *µ*(*i*))) is a diagonal weighted matrix, *µ* = [*µ*(1), *µ*(2),…] is the vector of expected values,

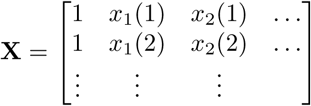

is the dataset in matrix form, and **y**(*i*) = [*y*(1), *y*(2),…]^*T*^ is the vector of response variables.

It can be observed that the **B** matrix is of dimensionality *N* × *N*, where *N* is the number of instances. For large volume of data, this matrix can greatly affect the performance of the implementation. However, we need to only keep the values along the diagonal as this is a diagonal matrix; thereby precluding the potential performance drawbacks. The pseudo-code of both glm and our implementation are given in Algorithm 1 and 2 respectively.

In our implementation, there are two hyper parameters that need to be tuned before running the process. One is the learning rate of the logistic regression model and the other is the number of maximum iterations allowed for convergence. We used maximum iteration as 25 because we found that the glm implementation of R has 25 maximum iteration by default [12]. We used the learning rate value of 0.1.

#### Algorithm 1 glm

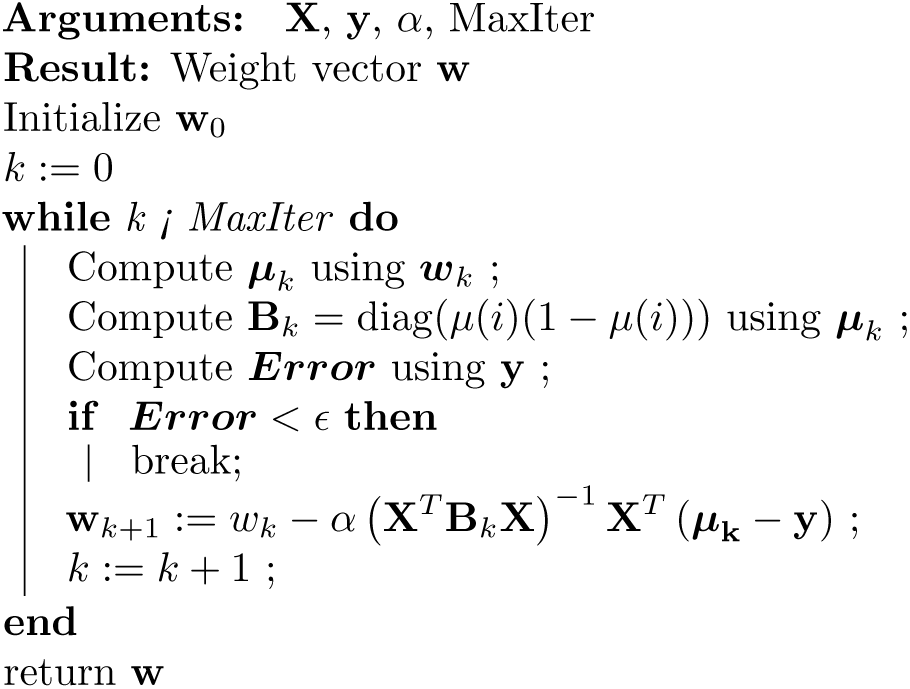

#### Algorithm 2 FastHawk

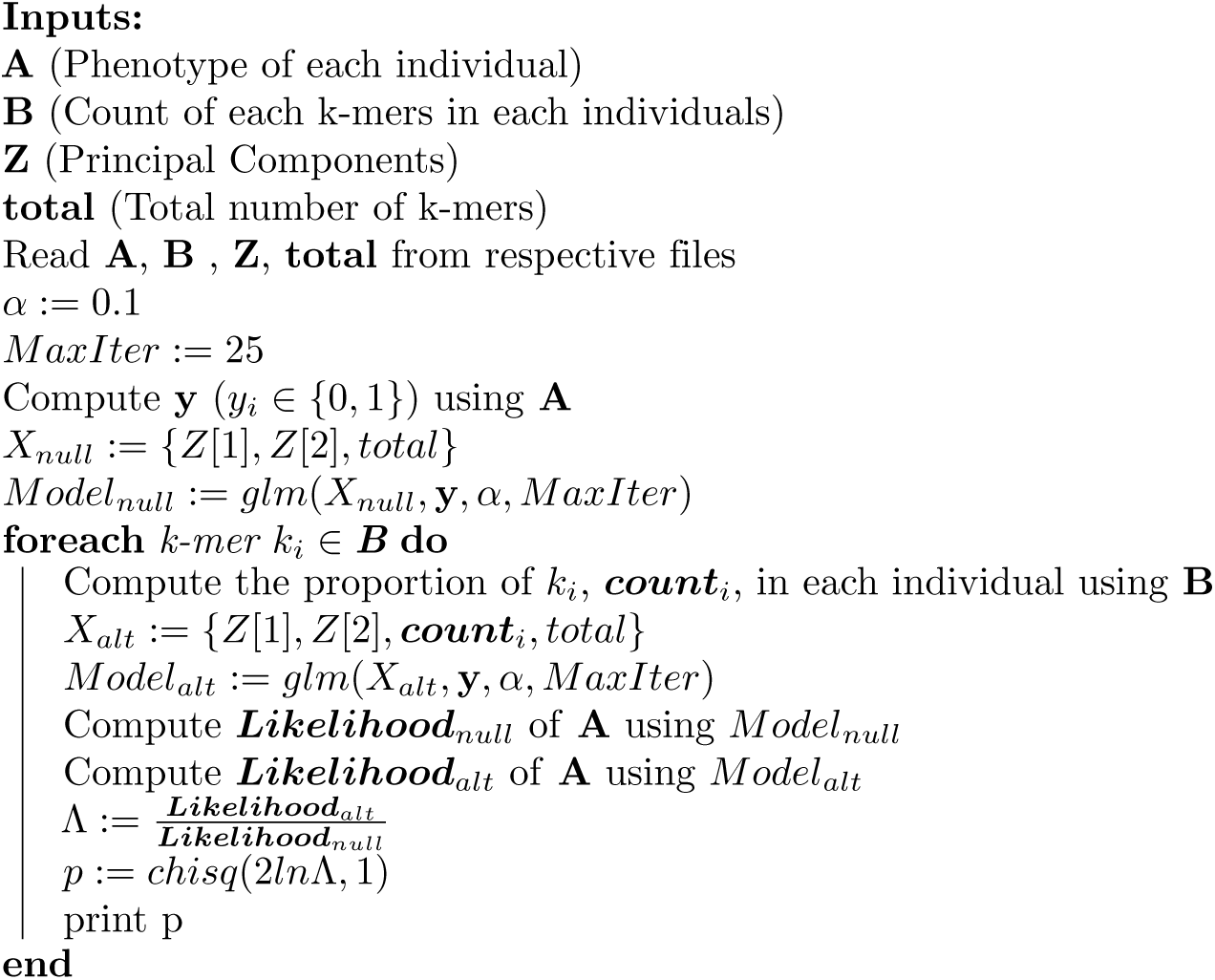

The implementation also makes it easier for users to specify number of principal components to be used for population structure correction as well as additional covariates using command line parameters and input files.

### Bug fixes

An error was found in the old implementation regarding the order of samples during adjustment of p-values using the confounding factors. This has been corrected in the new C++ implementation.

### Implementation of Benjamini–Hochberg Procedure

A number of corrections exist for p-value thresholds when multiple tests are being performed. Two such methods are: Bonferroni correction and Benjamini-Hochberg correction. The previous implementation performed hypothesis testing on each k-mer and performed Bonferroni correction using the total number of k-mers to determine k-mers associated with the phenotype in question. However, Bonferroni correction is known to be conservative i.e. it may fail to reject the null hypothesis even when it should be rejected.

Here, we implemented Benjamini-Hochberg correction which controls the false discovery rate (FDR). This can be used in studies underpowered for Bonferroni correction. The new implementations gives the provision for performing either correction according to user preference.

### Incorporation of Jellyfish 2

The original implementation of Hawk used a modified version of Jellyfish [13]. Subsequently, Jellyfish2 has been released which provides better performance. The present implementation of Hawk allows k-mer counting using a modified version of Jellyfish2 available through our Github repository.

### Re-implementation of post-processing

Once sequences corresponding to each loci associated with the phenotype are obtained, information such as average p-values of constituent k-mers as well as average number of times they are present in case and control samples could be looked up using scripts provided with the original implementation. However, these scripts used a combination of C++ codes and shell commands, and was found to be slow in some cases [5]. Here, we re-implement the script using hash tables in C++ to speed up the look-up.

## Results

To assess the performance and accuracy of our implementation, we use the *E*.*coli* dataset on ampicillin resistance which was analyzed using the original implementation of Hawk [1].

All the experiments are performed on a machine with CPU Intel(R) Xeon(R) CPU E5-2697 v2 @ 2.70GHz, 386GB memory with OS Ubuntu 18.04.3. There are two CPUs in the system with total 48 logical cores. The C++ implementation are compiled with g++-7.4.0 with no optimization flag. Execution time is measured using the “date” command.

### Hyper parameters

The code has 3 hyper parameters which are described below along with the values used in Table 1.

**Table 1.**
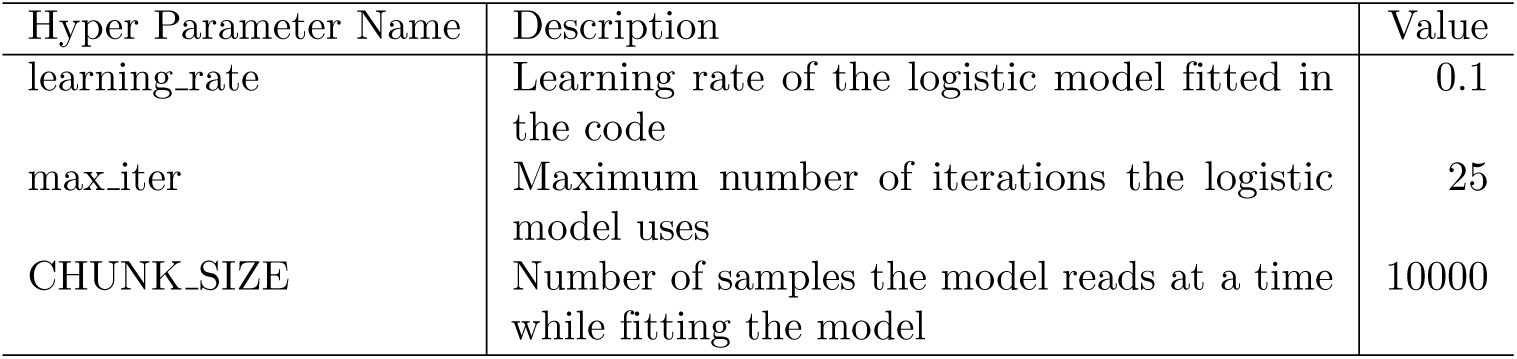
Hyper parameter values used in the implementation

| Hyper Parameter Name | Description | Value |
| --- | --- | --- |
| learning_rate | Learning rate of the logistic model fitted in the code | 0.1 |
| max_iter | Maximum number of iterations the logistic model uses | 25 |
| CHUNK_SIZE | Number of samples the model reads at a time while fitting the model | 10000 |

### Comparison of p-values

We have compared the output of our implementation and the output of the previous implementation by plotting logarithm of p-values obtained using the C++ implementation against that of p-values obtained using the R implementation in Figure 1. An identical result should give a line with slope 1. Graph produced by our implementation are almost linear with small deviation at few points.

**Figure 1.**
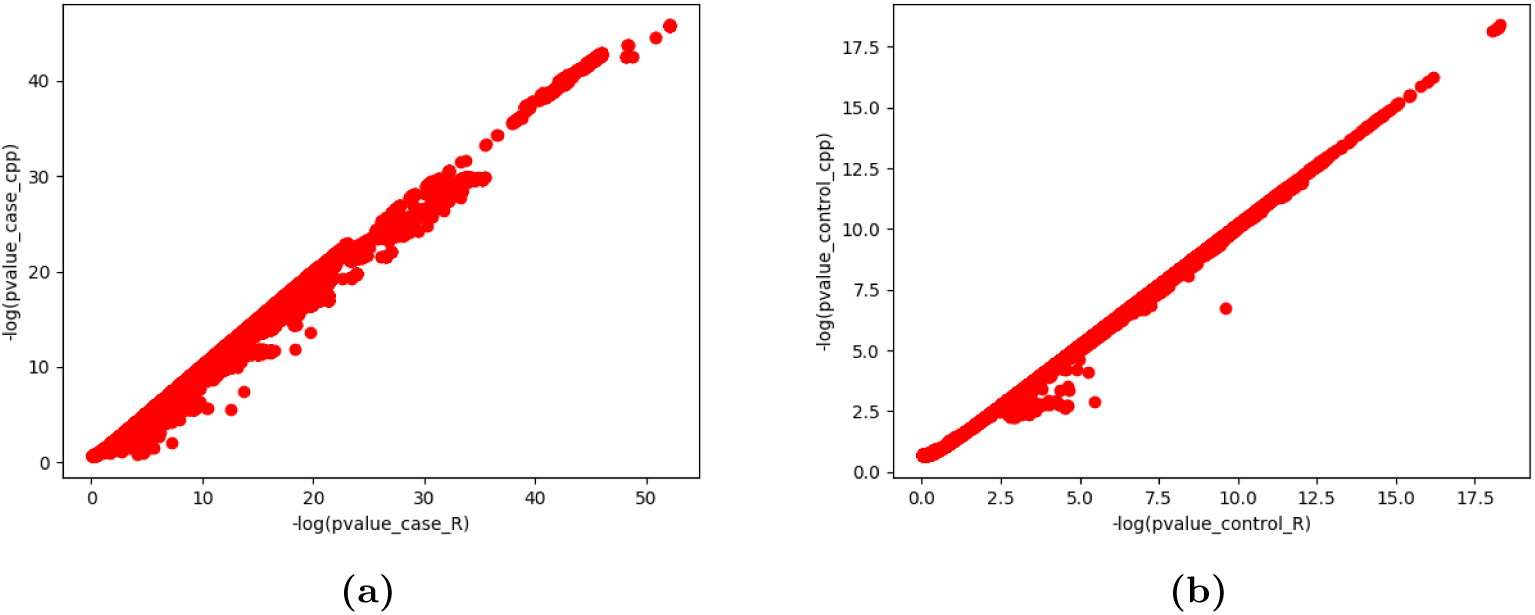
Comparison of log of p-values computed using the previous implementation in R and new implementation using C++ for k-mers of potential association with cases and controls

The k-mers found significant using the old implementation (corrected version) were mapped to *Escherichia coli* strain DTU-1 genome [GenBank: CP026612.1] and *Escherichia coli* strain KBN10P04869 plasmid pKBN10P04869A sequence [GenBank: CP026474.1]. The positions in the reference genomes and the p-values are shown in Manhattan plots in Figures 2(a) and (b). We find associations near the *β-lactamase TEM-1 (blaTEM-1)* gene, the presence of which is known to confer ampicillin resistance, as in [1]. However, some of the associations outside of this gene detected previously in [1], that are likely to be spurious, is no longer observed after the correction of error.

**Figure 2.**
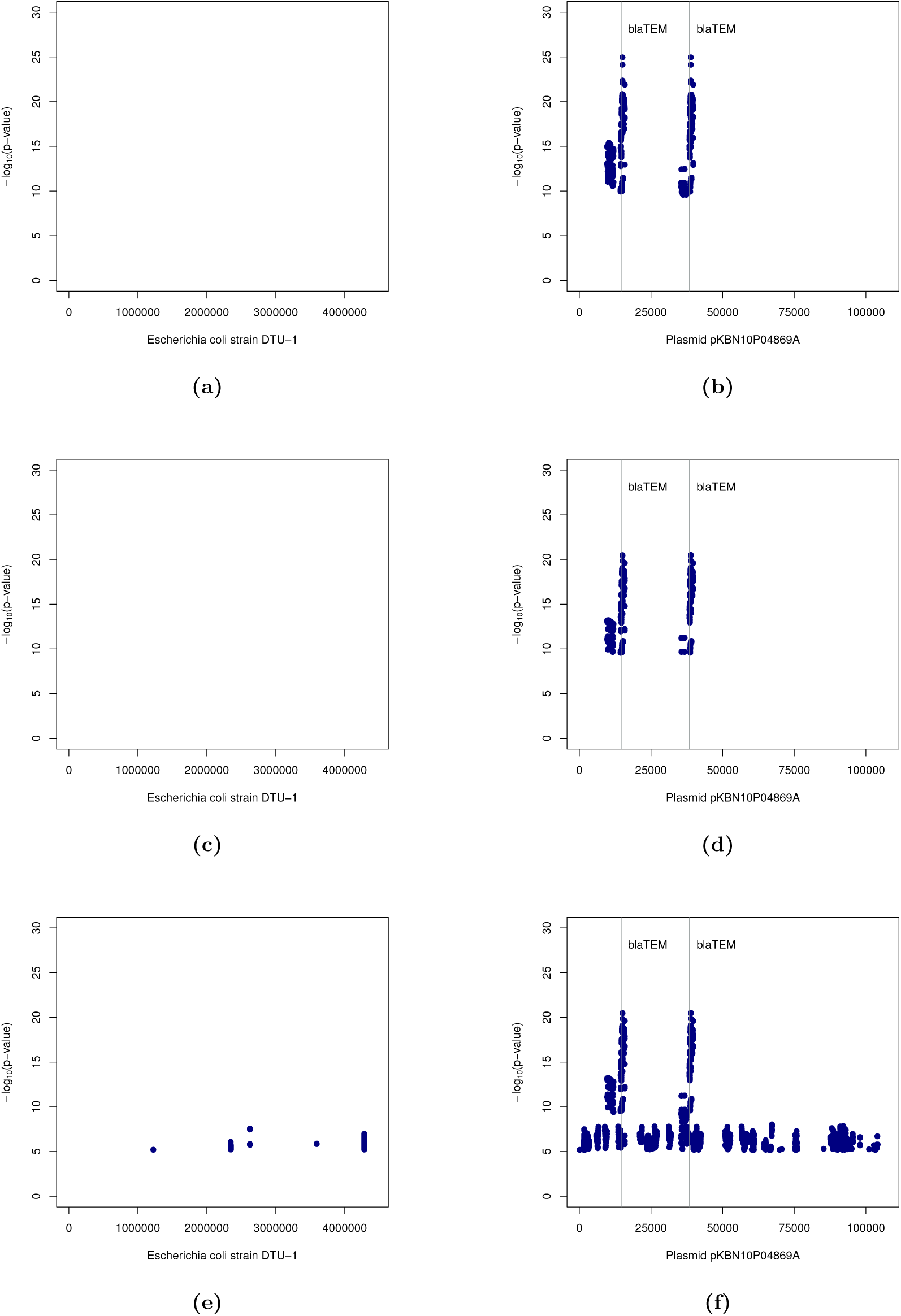
Manhattan plots showing negative logarithm of adjusted p-values of k-mers found significantly associated with ampicillin resistance against their start positions, computed using the R implementation with Bonferroni correction (a),(b); using C++ implementation with Bonferroni correction (c),(d); and using C++ implementation with Benjamini-Hochberg procedure (e),(f); in *Escherichia coli* strain DTU-1 genome (a), (c), (e), and plasmid pKBN10P04869A sequence (b), (d), (f). The vertical lines denote start positions of *β-lactamase TEM-1* gene, the presence of which is known to confer resistance to ampicillin.

The above analysis was also performed with the k-mers found significant using the new C++ implementation and the Manhattan plots are shown in Figures 2(c) and (d). We observe that associations are detected in same regions as those found using the R implementation.

### Controlling FDR using the Benjamini-Hochberg procedure

Hawk uses Bonferroni correction to address multiple testing by default. However, we provide the option to control false discovery rate (FDR) using the Benjamini-Hochberg procedure. Figures 2(e) and (f) show Manhattan plots when k-mers are considered associated with ampicillin resistance after controlling FDR using the Benjamini-Hochberg procedure. We observe that many k-mers outside of the *β-lactamase TEM-1* gene are considered significant. We therefore recommend using Bonferroni correction and using the Benjamini-Hochberg procedure only if the study is under-powered for Bonferroni correction.

### Comparison of running time

The execution of times of the old and new implementations are compared in Figure 3. Reported time is obtained by running both implementations using 32 threads. We find that the C++ implementation of the confounder correction phase is approximately three times faster than the precious implementation in R. However, the overall execution time is dominated by the k-mer counting step.

**Figure 3.**
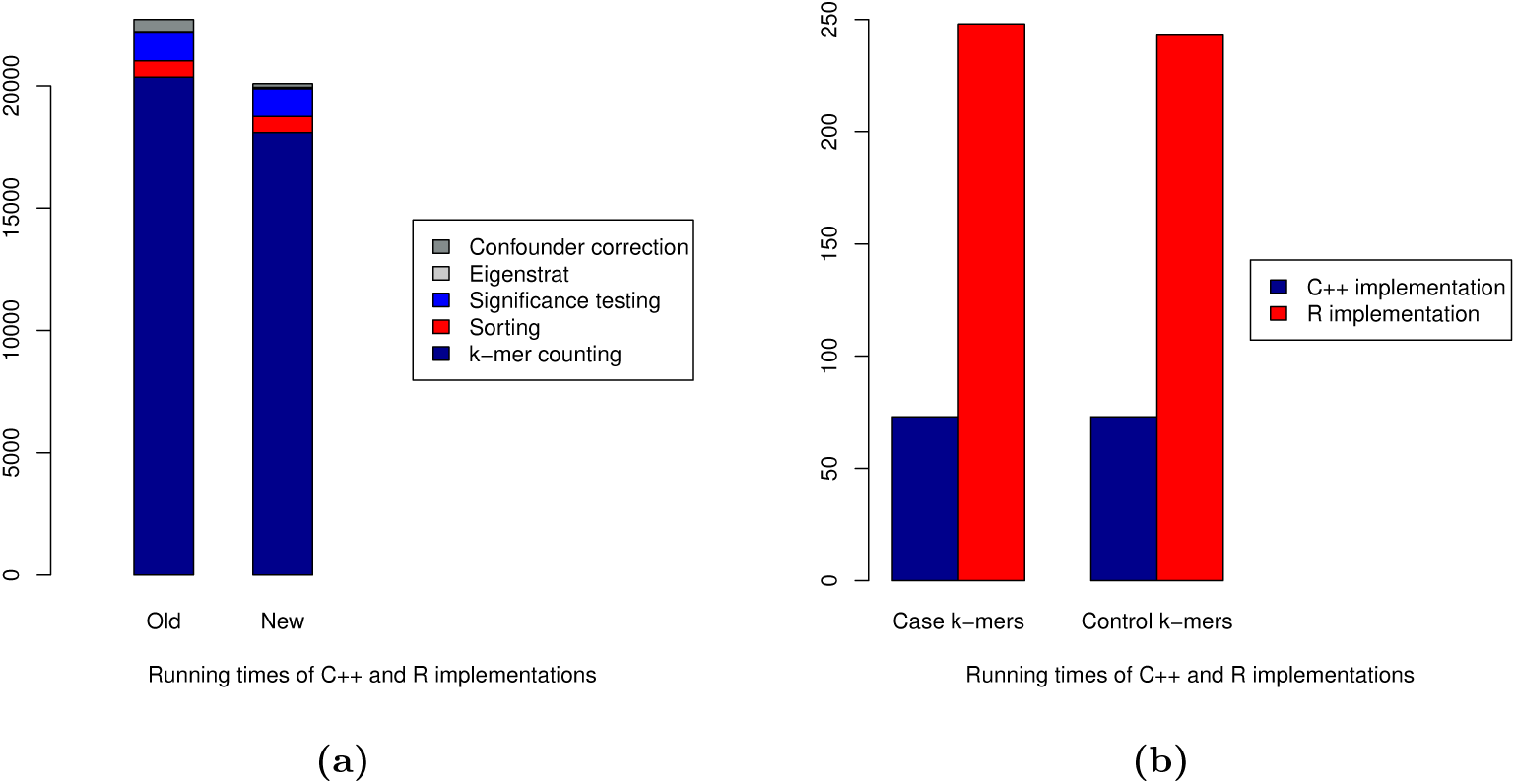
Comparison of execution times of old and new implementations (a) for the entire pipelines, and (b) correction of confounding factors using C++ and R implementations

Table 2 shows comparison of execution times of Jellyfish and Jellyfish 2. We observe that, although Jellyfish 2 is faster overall, the performance improvement it provides is not substantial.

**Table 2.** Comparison of running times of Jellyfish and Jellyfish 2

| Sub-command | Jellyfish (sec) | Jellyfish 2 (sec) |
| --- | --- | --- |
| histo | 20 | 674 |
| count | 19,852 | 16,580 |
| dump | 481 | 824 |
| Total | 20,353 | 18,078 |

### Finding sex-specific sequences

The HAWK pipeline can be used to find sequences of sex chromosomes in organisms with unassembled or poorly assembled genomes. To assess the performance, we ran HAWK on sequencing data from the Yoruba in Ibadan, Nigeria (YRI) and the Toscani in Italia (TSI) populations from the 1000 genomes project dataset [14]. Of the 110 YRI and 109 TSI individuals, 107 were male and 112 were female. The sexes of individuals were used as cases and controls and the pipeline was executed.

The initial step revealed 106,272,845 and 17,056,781 k-mers that are present significantly more times in female and male samples respectively compared to the other. Principal component analysis was then performed on the binary matrix denoting presence or absence of 32,699,548 randomly chosen k-mers, present between 1% and 99% of the samples. The PCA plots are shown in Figure 4. We observe that the first two PCs capture the population structure and hence they are used as confounders, as done by default. It may be noted that the fifth and sixth PCs nearly separate the two sexes. Treating these as confounders would lead to removal of many sex-specific k-mers.

**Figure 4.**
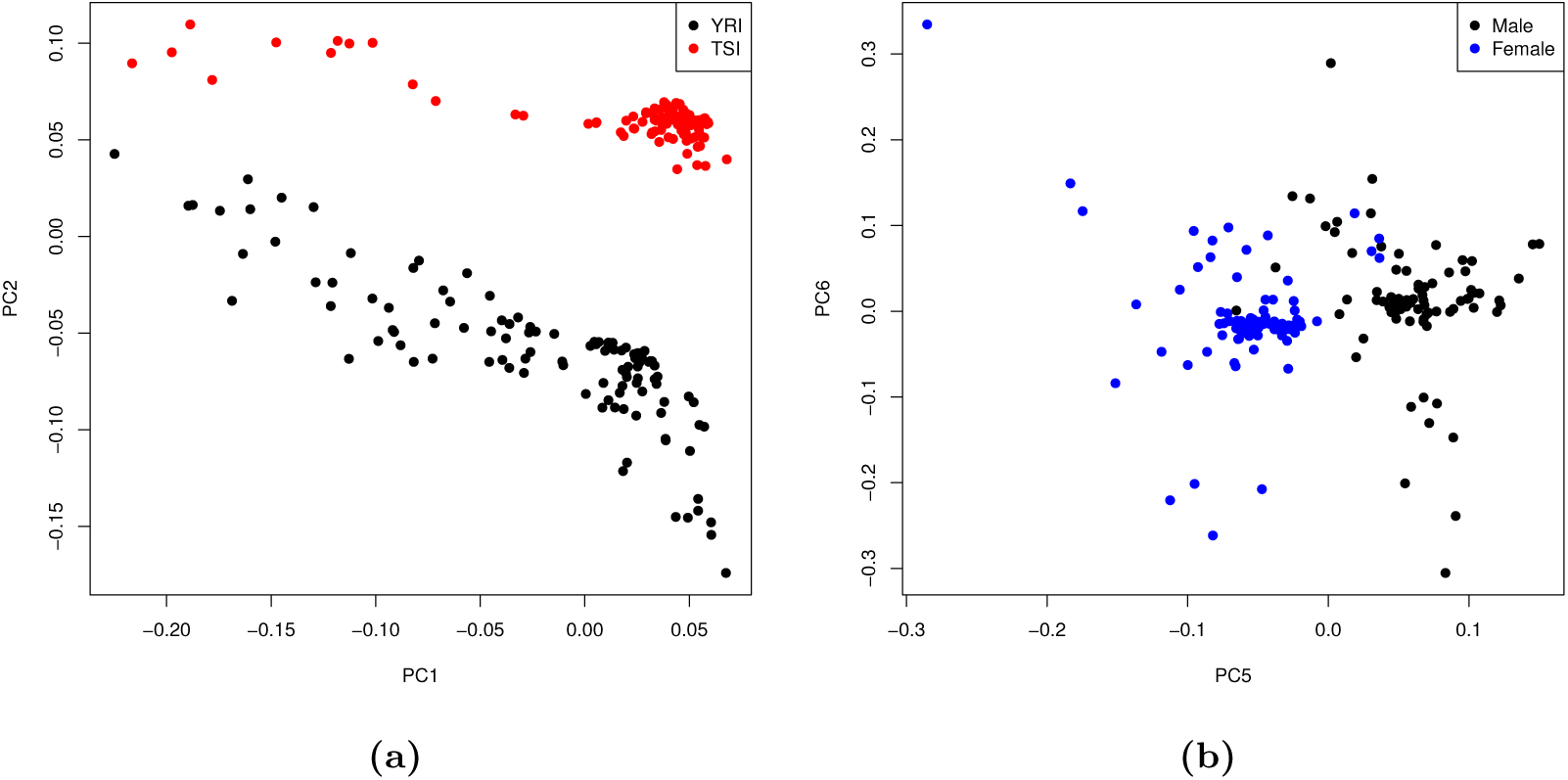
Principal component analysis (PCA) plots of the samples in the space formed by (a) first and second principal components with samples colored according to populations, and (b) fifth and sixth principal components with samples colored according to sex

After correcting for confounders, we obtain 14,473,058 and 54,256,206 k-mers associated with male and female samples respectively. The k-mers were mapped to the human reference genome using Bowtie 2 [15] to analyze their locations. The results are summarized in Table 3. We find that 99.97% and 96.37% of k-mers associated with female and males samples map to Chromosome X and Chromosome Y respectively. The remaining k-mers map to other locations in the human reference genome or stay unmapped.

**Table 3.** Summary of k-mers found associated with female and male samples

| Sex | Total | Chr X | Chr Y | Others | Unmapped |
| --- | --- | --- | --- | --- | --- |
| Female | 54,256,206 | 54,241,253 | 148 | 1,474 | 13,331 |
| Male | 14,473,058 | 6,197 | 13,947,961 | 63,454 | 455,446 |

The positions of the k-mers, associated with female and male samples, in Chromosomes X and Y respectively are shown in Figure 5. We observe that k-mers throughout the entire sequenced regions of the two chromosomes are detected using HAWK. We note that the region in Chromosome Y, where no k-mer could be mapped, is missing from the reference genome [16]. The missing region in Chromosome Y also possibly explains the large percentage of male associated k-mers that could not be mapped to it in comparison to the percentage of female associated k-mers that map to Chromosome X.

**Figure 5.**
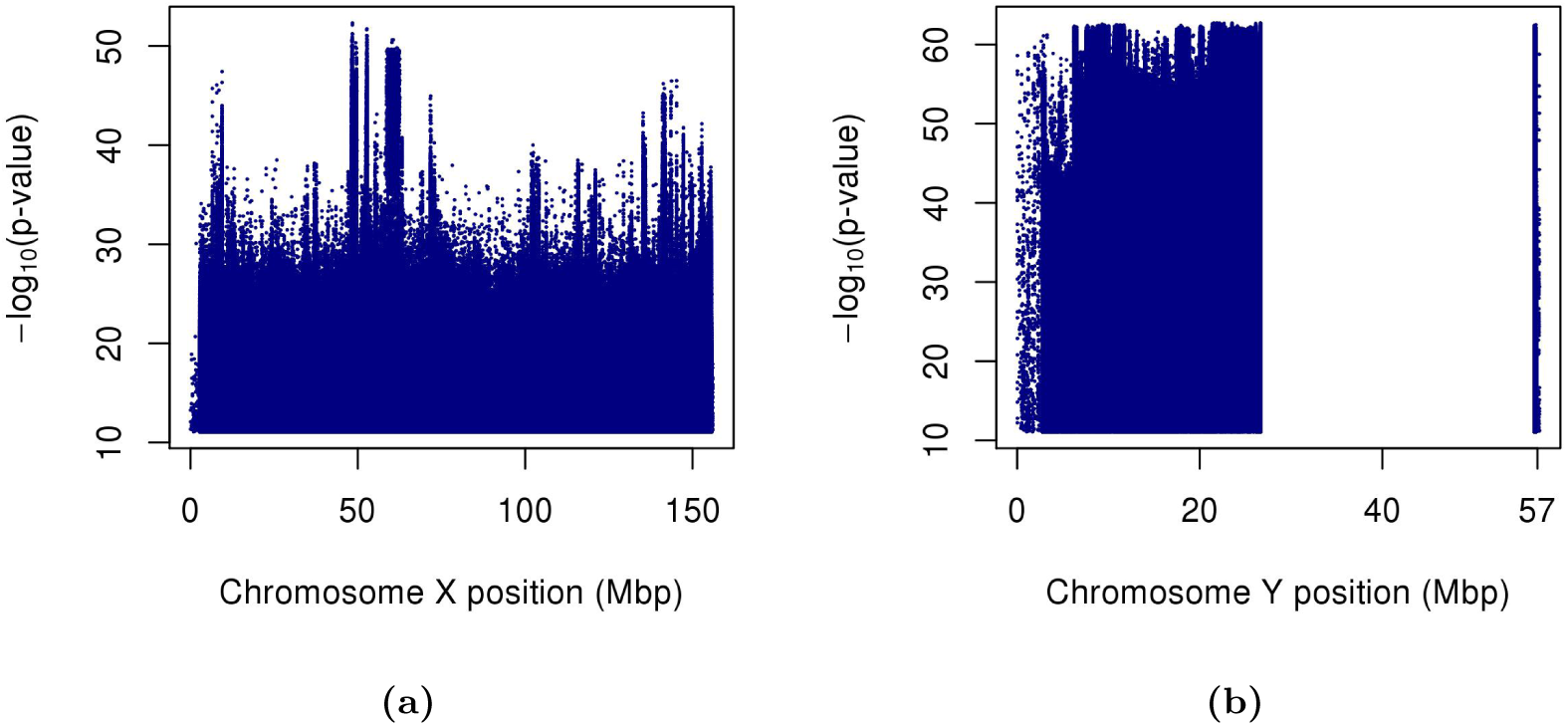
Manhattan plots showing negative logarithm of p-values of (a) k-mers associated with female samples against their positions in Chromosome X, and (b) k-mers associated with male samples against their positions in Chromosome Y. The region in Chromosome Y where no k-mer mapped is missing from the reference genome

## Conclusions

We have re-implemented portions of Hawk, which is a tool for association mapping using k-mers. The re-implementation in C++ makes it faster and more convenient to use while retaining accuracy. We have also added support for the new version of k-mer counting tool Jellyfish and correction for multiple testing using the Benjamini-Hochberg procedure. The k-mer counting step remains the bottleneck in the pipeline which may be addressed by adding support for other k-mer counting tool. Finally, we show how the method can be applied to determine sex specific sequences in organisms accurately.

## Availability and User Support

Scripts, source code, and user manual for Hawk is available at https://github.com/atifrahman/HAWK. The system is available for and tested in Linux (Ubuntu14.04 and Ubuntu18.04).

## Acknowledgments

We like to thank Guillaume Marcais for providing the patch to modify Jellyfish 2 [13] according to the requirements of HAWK.

## References

1. Atif Rahman, Ingileif Hallgrímsdóttir, Michael Eisen, and Lior Pachter. Association mapping from sequencing reads using k-mers. eLife, 7:e32920, 2018.

2. Samuel K Sheppard, Xavier Didelot, Guillaume Meric, Alicia Torralbo, Keith A Jolley, David J Kelly, Stephen D Bentley, Martin CJ Maiden, Julian Parkhill, and Daniel Falush. Genome-wide association study identifies vitamin b5 biosynthesis as a host specificity factor in campylobacter. Proceedings of the national academy of sciences, 110(29):11923–11927, 2013.

3. John A Lees, Minna Vehkala, Niko Välimäki, Simon R Harris, Claire Chewapreecha, Nicholas J Croucher, Pekka Marttinen, Mark R Davies, Andrew C Steer, Steven YC Tong, et al. Sequence element enrichment analysis to determine the genetic basis of bacterial phenotypes. Nature communications, 7:12797, 2016.

4. Sarah G Earle, Chieh-Hsi Wu, Jane Charlesworth, Nicole Stoesser, N Claire Gordon, Timothy M Walker, Chris CA Spencer, Zamin Iqbal, David A Clifton, Katie L Hopkins, et al. Identifying lineage effects when controlling for population structure improves power in bacterial association studies. Nature microbiology, 1:16041, 2016.

5. Magali Jaillard, Leandro Lima, Maud Tournoud, Pierre Mahé, Alex Van Belkum, Vincent Lacroix, and Laurent Jacob. A fast and agnostic method for bacterial genome-wide association studies: Bridging the gap between k-mers and genetic events. PloS genetics, 14(11):e1007758, 2018.

6. Yoav Voichek and Detlef Weigel. Identifying genetic variants underlying phenotypic variation in plants without complete genomes. Nature Genetics, 2020.

7. Guillaume Marçais and Carl Kingsford. A fast, lock-free approach for efficient parallel counting of occurrences of k-mers. Bioinformatics, 27(6):764–770, 2011. doi: 10.1093/bioinformatics/btr011. URL http://bioinformatics.oxfordjournals.org/content/27/6/764.abstract.

8. Nick Patterson, Alkes L Price, and David Reich. Population structure and eigenanalysis. PLoS genetics, 2(12):e190, 2006.

9. Alkes L Price, Nick J Patterson, Robert M Plenge, Michael E Weinblatt, Nancy A Shadick, and David Reich. Principal components analysis corrects for stratification in genome-wide association studies. Nature genetics, 38(8):904, 2006.

10. Yoav Benjamini and Yosef Hochberg. Controlling the false discovery rate: a practical and powerful approach to multiple testing. Journal of the royal statistical society. Series B (Methodological), pages 289–300, 1995.

11. R manual. Fitting Generalized Linear Models. URL https://stat.ethz.ch/R-manual/R-devel/library/stats/html/glm.html.

12. DataScience StackExchange. Number of Iterations in R glm. URL https://datascience.stackexchange.com/a/16811.

13. Guillaume Marçais and Carl Kingsford. A fast, lock-free approach for efficient parallel counting of occurrences of k-mers. Bioinformatics, 27(6):764–770, 2011.

14. Genomes Project Consortium et al. An integrated map of genetic variation from 1,092 human genomes. Nature, 491(7422):56–65, 2012.

15. Ben Langmead and Steven L Salzberg. Fast gapped-read alignment with bowtie 2. Nature methods, 9(4):357, 2012.

16. Miten Jain, Hugh E Olsen, Daniel J Turner, David Stoddart, Kira V Bulazel, Benedict Paten, David Haussler, Huntington F Willard, Mark Akeson, and Karen H Miga. Linear assembly of a human centromere on the y chromosome. Nature biotechnology, 36(4):321–323, 2018.

